# Systematic review and meta-analysis of protocols and yield of direct from sputum sequencing of *Mycobacterium tuberculosis*

**DOI:** 10.1101/2024.12.04.625621

**Authors:** B.C. Mann, J. Loubser, S. Omar, C. Glanz, Y. Ektefaie, K.R. Jacobson, R.M. Warren, M.R. Farhat

## Abstract

Direct sputum whole genome sequencing (dsWGS) can revolutionize *Mycobacterium tuberculosis* (*Mtb*) diagnosis by enabling rapid detection of drug resistance and strain diversity without the biohazard of culture. We searched PubMed, Web of Science and Google scholar, and identified 8 studies that met inclusion criteria for testing protocols for dsWGS. Utilising meta-regression we identify several key factors positively associated with dsWGS success, including higher *Mtb* bacillary load, mechanical disruption, and enzymatic/chemical lysis. Specifically, smear grades of 3+ (OR = 14.7, 95% CI: 3.5, 62.1; p = 0.0005) were strongly associated with improved outcomes, whereas decontamination with sodium hydroxide (NaOH) was negatively associated (OR = 0.005, 95% CI: 0.001, 0.03; p = 7e-06), likely due to its harsh effects on *Mtb* cells. Furthermore, mechanical lysis (OR = 193.3, 95% CI: 11.7, 3197.8; p = 0.008) and enzymatic/chemical lysis (OR = 18.5, 95% CI: 1.9, 183.1; p = 0.02) were also strongly associated with improved dsWGS. Across the studies, we observed a high degree of variability in approaches to sputum pre-processing prior to dsWGS highlighting the need for standardized best practices. In particular we conclude that optimizing pre-processing steps including decontamination with the exploration of alternatives to NaOH to better preserve Mtb cells and DNA, and best practices for cell lysis during DNA extraction as priorities. Further and considering the strong association between *Mtb* load and successful dsWGS, protocol improvements for optimal sputum sample collection, handling, and storage could also further enhance the success rate of dsWGS.

## Introduction

Tuberculosis (TB) caused by the bacillus *Mycobacterium tuberculosis* (*Mtb*) remains the most common cause of death from any single infectious pathogen (1, 2). Progress in eradicating TB is hampered by the emergence of multidrug-resistant (MDR) and extensively drug resistant (XDR) *Mtb* strains. According to the latest WHO Global Tuberculosis Report approximately half a million people worldwide developed rifampicin-resistant TB (RR-TB), 78% of which had MDR-TB, defined as resistant to at least isoniazid (INH) and rifampicin (RIF) (2). Next Generation Sequencing (NGS) advances over the past decade have provided the ability to rapidly sequence the whole *Mycobacterium tuberculosis* genome; supplying an extraordinary tool to study the genetic epidemiology of this pathogen, while also detecting single nucleotide polymorphisms (SNPs) and other mutations which can be used to predict susceptibility to first-line drugs (3, 4). Several rapid genotypic drug susceptibility testing (DST) methods are currently endorsed by the WHO, but most of these tests do not provide a comprehensive summary of a patient’s drug resistance (DR) profile, since most of these assays only focus on a limited number of targets involved with DR (5). Recently the introduction of targeted NGS (tNGS) has expanded the number of drugs and targets assayed substantially (6). Whole genome sequencing (WGS) however can assay the full breadth of genetic variation and is most comprehensive for predicting phenotype from genotype for *Mtb* (7, 8).

*Mtb* sequencing is still hindered by the long and cumbersome process of culturing *Mtb* for DNA extraction. This process can take weeks and sometimes even months (4). In addition to the long culture period, the culture-based approach has an additional limitation, as culture can change the population structure of the original sample due to the selection of subpopulations more suited to growth in culture and random genetic drift (9).The logical step to avoid these limitations is to sequence *Mtb* directly from clinical specimens. To date several studies have demonstrated that sequencing from direct patient specimens is possible with varying levels of success (3, 4, 9–12). Commercial tNGS assays include a targeted polymerase chain reaction (PCR) amplification step that helps improve sequencing coverage from direct samples, however even tNGS performs poorly on sputum with lower bacillary burden (including Xpert medium and low, or smear negative sputum) with 50-78% or more of test samples yielding only partial coverage of the targets (6, 8, 13). For direct whole genome sequencing, most studies have relied on the use of target capture and enrichment technology which enriches target DNA with a set of target-specific bait probes during library preparation. Others have opted for a selective lysis approach, attempting to selectively lyse contaminating host and bacterial cells, followed by the depletion of contaminating DNA by enzymes such as DNase, either forgoing or only turning to target enrichment in samples for which it was deemed necessary by additional quality control (QC) steps (3, 4, 10, 11). This systematic review and meta-analysis aims to summarize DNA target and non-target-based methods previously utilized to perform whole genome sequencing *Mtb* from direct patient samples. We classify the different approaches used in the literature and perform an individual sample meta-analysis of the effect of specific sample processing steps on direct sequencing success. Although focused on WGS, reviewing non-target-based methods can also have implications for improving yield of tNGS from direct patient samples.

## Methods

### Search strategy

In brief we searched PubMed and Web of Science, looking for English articles published on *Mtb* direct sputum whole genome sequencing (dsWGS) to compare various approaches that have previously been applied for successful dsWGS of *Mtb*. This was done to highlight gaps in the current applied methodology that can be improved upon to better facilitate dsWGS. The literature search was carried out using the following keywords “direct + sputum + sequencing + tuberculosis + mycobacteria” and was conducted by three independent reviewers. The three reviewers reviewed titles, abstract, key words and subsequently the full texts to identify and include articles meeting the study inclusion and not meeting the exclusion criteria below.

An ethical review was deemed unnecessary as this was a secondary analysis of published articles. We conducted an additional search of Google scholar to identify any relevant articles not identified in the original search. No grey literature, conference papers or unpublished works were included in this analysis because of uncertainty over the relevance and validity of the presented data.

### Inclusion and exclusion criteria

The inclusion criteria for eligible publications were defined as all articles that:

- Studied *Mtb*.
- Attempted direct whole genome sequencing or target capture/target enrichment followed by whole genome sequencing (i.e. no targeted PCR) directly on sample without intervening culture.
- Reported on *Mtb* input sputum smear grade or Xpert CT or other *MTB* input DNA quantification.
- Reported on one or more of the following outputs:

`o` *Mtb* genomic coverage of drug resistance genes (any subset) or the whole genome.
`o` Resistance mutation recall relative to phenotype or to sequencing after culture.
`o` Lineage mutation recall relative to sequencing after culture.
- Provided methodological detail on the sputum processing protocol.

The exclusion criteria included articles that:

- Focused on the application to pathogens other than *Mtb*.
- Focused on samples other than sputum.

### Data extraction

Three reviewers independently extracted data from the included studies.The following background information was extracted from eligible papers: Author details, year of publication and population. Sixteen technical variables were extracted under three categories including Sample data and characteristics, Methodology and Results (Table 1). Data extracted were compiled into several predesigned spreadsheets. In instances where data was represented graphically, the authors were contacted to provide the original numerical data used to generate the graphs. If the authors could not be reached or the information was no longer available, we extracted the numerical values from the graphical representations using available software (PlotDigitizer).

**Table 1.** Summary of data items extracted from articles fulfilling inclusion and exclusion criteria.

| Data group | Number | Data extracted |
| --- | --- | --- |
| **Sample data and characteristics** | 1 | Population group |
|  | 2 | Sample number |
|  | 3 | Sample types |
|  | 4 | Sputum culture pairs |
|  | 5 | Xpert or qPCR data (quantitative or semiquantitative) |
|  | 6 | Smear data |
| **Methodology** | 7 | Sample pre-treatment and enrichment methodology |
|  | 8 | DNA extraction methodology |
|  | 9 | DNA concentrations |
|  | 10 | Bioinformatics methodology |
| **Results** | 11 | Coverage for both culture and direct sputum samples |
|  | 12 | Percentage on target reads for direct sputum samples |

### Bioinformatics analysis

Sequencing data was downloaded for each included study, either from NCBI GenBank or the European Nucleotide Archive (ENA). Initial quality assessment was done using fastQC, version 0.11.9, followed by adapter removal, quality filtering and per-read low quality base trimming using fastP, version 0.20.1 (14). Following quality control, reads were taxonomically classified using the metagenomic classification tool Kraken2 using the standard database (version 2.0.8) (Wood *et al*., 2019). Mycobacterial reads were extracted and aligned using bwa-mem2 (version 2.2.1) to the H37Rv reference genome AL123456 (15). Duplicates were removed using Picard and excluded in downstream analyses. Alignment statistics, including number of reads, depth and breadth of coverage and GC-content were determined and visualised using Qualimap (version 2.2.2c) (16).

### Statistical analysis

To assess the effects of various factors on genome and DR region coverage (DR regions comprised of 73 genomic regions strongly associated with a DR phenotyope, identified and curated by the well-known TB-Profiler tool) (17), we employed regression comparing generalized linear mixed models (GLMMs) to support batch control across studies, and a generalized linear model (GLM) without batch control. We found no significant batch effects across the studies and report the GLM results in the main text and the GLMM results in the supplement (Supplementary Tables Y and Z). We assessed the fixed effects of several factors including: smear grade, mechanical disruption, enzymatic/chemical lysis, decontamination, and heat treatment on whole genome coverage (>5x at >90%) and coverage of DR conferring regions (>5x at >95%). Given the limited number of samples sequenced directly we limited the analysis to samples that underwent target capture and enrichment and excluded directly sequenced samples. Associations were assessed using the Wald test with significance assessed at a P-value <0.05. Two processing steps homogenisation and contamination depletion were coded but excluded from the final model due to their utilization in all but one study, or only two target capture studies respectively making it difficult to evaluate their effect.

## Results

### Search strategy and study selection

We identified 134 and 91 studies respectively from the PubMed and Web of Science searches. One additional record was identified using a Google scholar search. Three independent reviewers read titles, abstracts, and keywords to assess for duplicates and adherence to inclusion and exclusion criteria. Records that were not excluded at this stage were reviewed in full text to assess the same; 219 records were excluded leaving a total of 8 studies for inclusion in this review.

### Characteristics of included studies

The number of participants per study ranged from 34 to 100; 5 studies sequenced directly from sputum and compared with sequencing after culture. The largest number of sputum/ culture pairs available from a single study was 43. All studies except 1 included smear data as a semi quantitative measure of bacilli load, and 3 studies included Xpert cycle threshold (Ct) values. Only 2 studies quantified DNA in the input sample by means of qPCR, an additional 2 studies report that qPCR was done to assess bacillary load, but results were not reported, or accessible after contacting the authors (Supplementary Table 1).

### Current approaches facilitating direct sputum sequencing from clinical specimens

The selected studies spanned three primary avenues to facilitate dsWGS with various ways of pre-treating samples before proceeding to one or the other (Figure 2). We define these methods broadly as: 1) non-(DNA) target methods: including selective lysis or other physical or chemical enrichment, which involves the enrichment of *Mtb* cells by breaking down contaminating host and/or commensal bacterial cells, followed by depletion of contaminating DNA by washing or enzymatic degradation and then sequencing; 2) DNA targeting methods specifically the bait capture approach that capture and enrich for *Mtb* DNA with specific DNA/RNA bait probes and; 3) a combination approach involving both selective lysis and contaminating DNA depletion, while also employing bait capture probes (3, 4, 9–12, 18).

**Figure 1:**
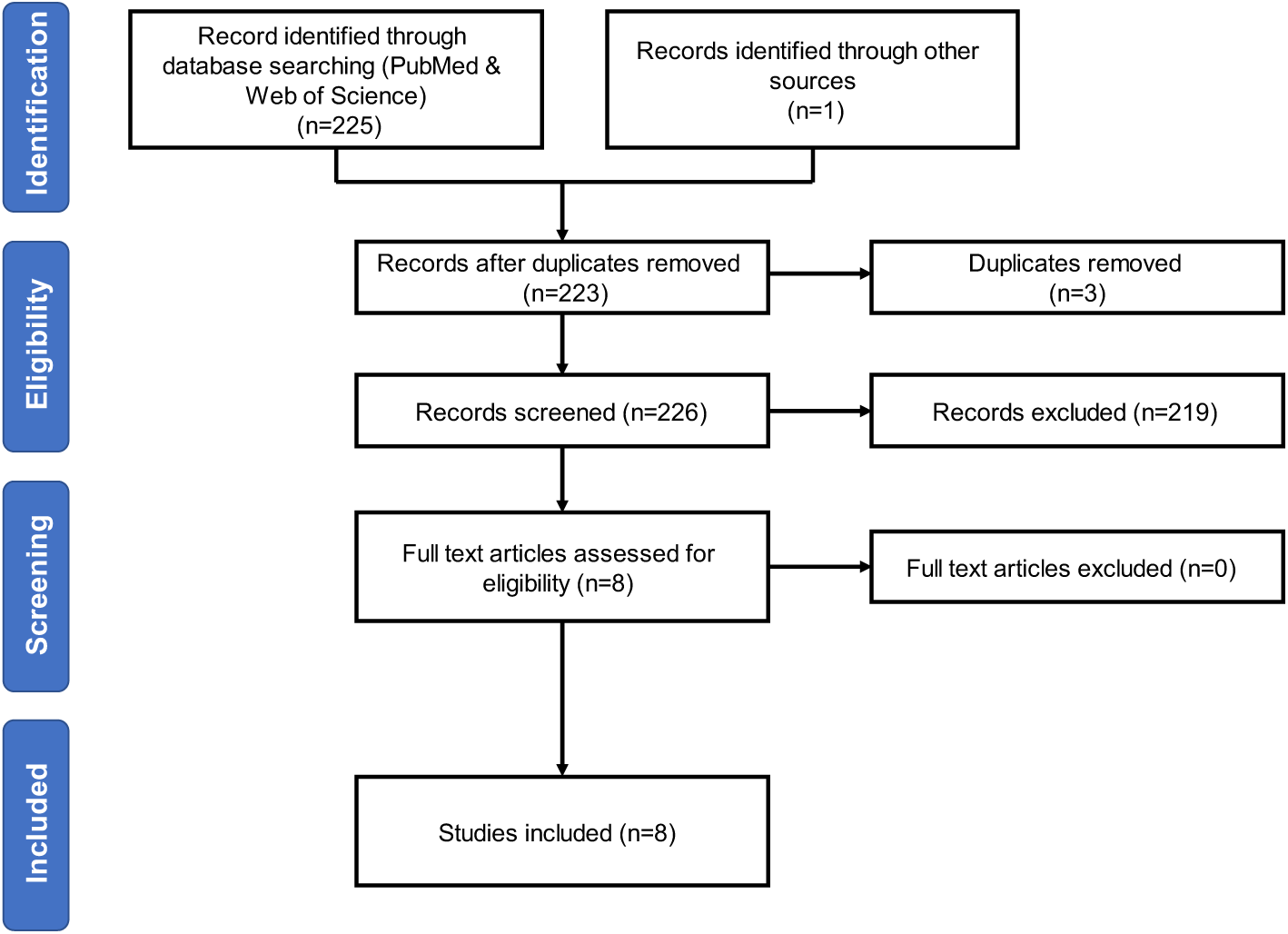
Flow diagram outlining study selection. Abbreviation: PRISMA, Preferred Reporting Items for Systematic Reviews and Meta-Analyses.

**Figure 2:**
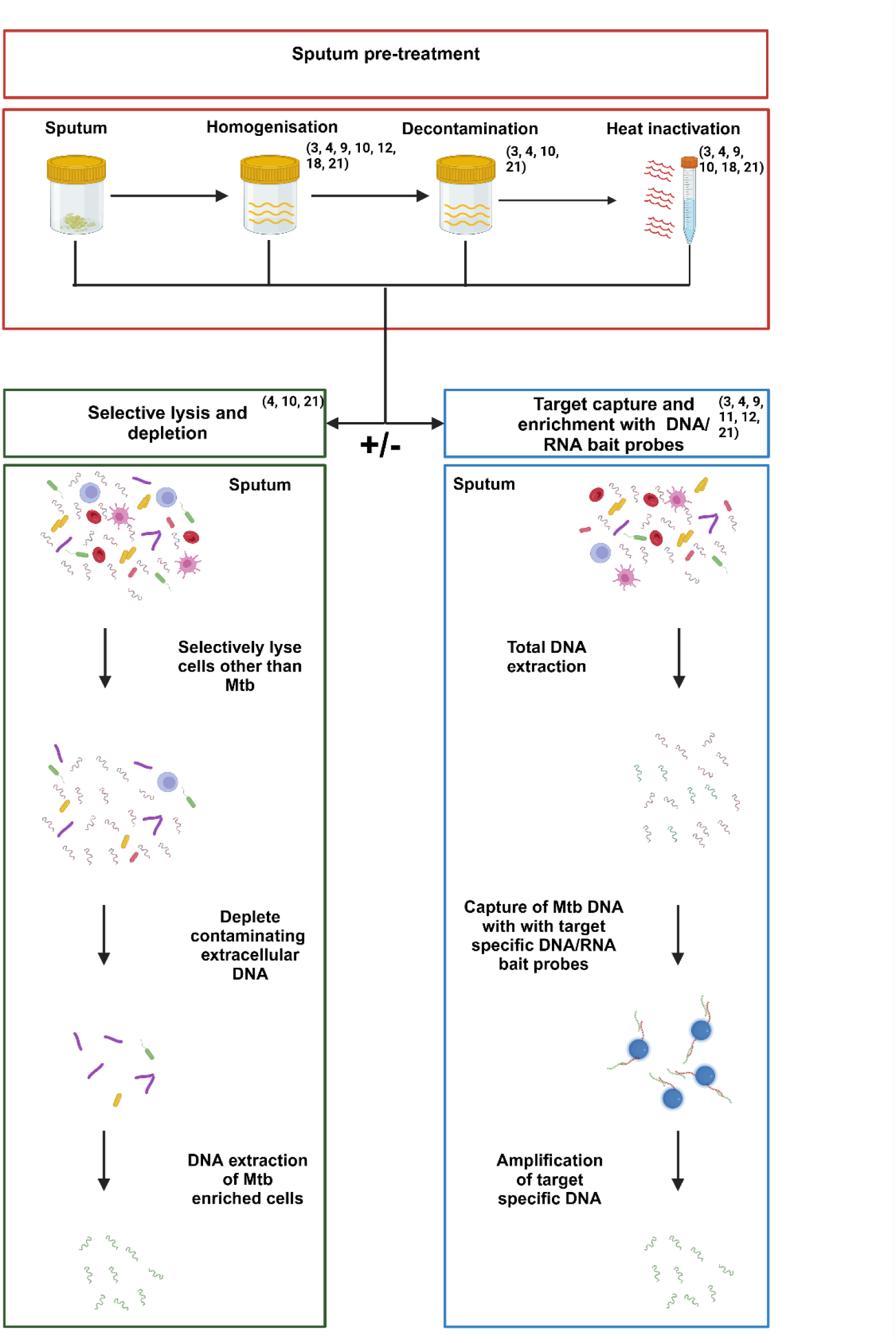
Illustration of potential pre-treatment steps that may be applied to sputum samples, prior to direct sequencing, while also demonstrating the alternative approaches that may follow any of the pre-treatment steps. This includes 1) the selective lysis approach for further *Mtb* enrichment where non *Mtb* cells and their DNA are depleted by chemical and/or enzymatic means, or 2) the target capture and enrichment approach where target specific DNA is enriched by means of DNA/RNA bait probes. Studies numbered in the figure as they appear in the bibliography.Created in https://BioRender.com

### Key pre-processing steps that may contribute to the success of dsWGS

We aimed to identify key steps that may contribute to the enrichment of the *Mtb* target and thus success of dsWGS (Table 2). A step was marked with X if it formed part of the workflow described in the paper. We identified seven steps within either a targeted or non-targeted approach that were employed across the eight studies: specifically, sputum homogenisation, sputum decontamination, heat inactivation, host DNA depletion, commensal microbe DNA depletion, DNA extraction and target enrichment. We briefly summarize the approaches taken under each step below. A more detailed summary of the methodology applied at each step can be found in Supplementary Table 2.

**Table 2.** Direct sputum processing workflow as outlined for each individual study.

| Study (reference) | Homogenisation | Decontamination | Heat inactivation | Host DNA depletion | Depletion of commensal microflora | DNA extraction |  |  | DNA/RNA bait capture |
| --- | --- | --- | --- | --- | --- | --- | --- | --- | --- |
|  |  |  |  |  |  | Mechanical | Enzymatic | Chemical |  |
| Brown *et al.* 2015 (3) | X | X | X |  |  | X |  |  | X |
| Votintseva *et al.* 2017 (10) | X | X | X | X |  | X |  |  |  |
| Doyle *et al.* 2018 (11) |  |  |  |  |  | X | X | X | X |
| Nimmo *et al.* 2019 (9) | X |  | X |  |  |  | X | X | X |
| Soundararajan *et al.* 2020 (12) | X |  |  | X |  | X | X | X | X |
| Goig *et al.* 2020 (4) | X | X | X | X | X | X |  |  | X |
| George *et al.* 2020 (18) | X |  | X |  |  | X |  |  |  |
| Macedo *et al.* 2023 (21) | X | X | X |  |  |  | X | X | X |

### Sputum homogenisation and decontamination

Although none of the studies utilised homogenisation or decontamination with the aim of enriching for the *Mtb* target, these steps were still incorporated as part of sample pre-processing prior to DNA extraction. All studies reviewed, except one, included a homogenisation step, but approaches varied from the use of the mucolytic agent N-acetyl-L-cysteine (NALC) to the reducing agent dithiothreitol (DTT) or other DTT containing products such as Sputasol. After homogenisation sputum samples are decontaminated in four studies using sodium hydroxide (NaOH) with the goal of preferentially killing other bacteria, fungi, and viruses, thereby reducing the risk of these contaminants influencing diagnostic testing (Supplementary Table 3).

### Heat inactivation

All the studies reviewed except two applied a heat inactivation step commonly used to reduce the biohazard involved with downstream processing (Table 2 and 3). Heat inactivation of *Mtb* specimens involved exposing samples to high temperatures ranging between 80 and 95°C for times ranging from 15 min to 1 hour defined period (3, 4, 9, 11, 12, 18, 21). One of the studies (George *et al*. 2020) demonstrated that heat inactivation can achieve enrichment of tough to lyse cells such *Mtb* but required the addition of a specialised thermal protection buffer to maintain the integrity of *Mtb* DNA during extensive heating (30 min at 99°C) which also subsequently lead to the degradation of any extracellular host DNA (18).

### Host and commensal microbe DNA depletion

Votinseva *et al*. (2017) and Soundararajan *et al*. (2020), aimed to enrich for *Mtb* by applying commercially available kits namely the MolYsis Basic5 kit and the Ultra Deep Microprep DNA isolation kit (Molzym, Germany) for the depletion of host DNA prior to *Mtb* WGS. Goig *et al*. (2020), used GTC solution (4M guanidinium thiocyanate 4M, 0·5% w/v sodium N-lauryl sarcosine, 25mM trisodium citrate, 0·1M 2mercaptoethanol, 0·5% w/v Tween 80) instead to lyse eukaryotic cells in conjunction with DNase. In addition to this, the study by Galo *et al*. (2020), was also the only study to directly preform additional depletion with the aim to not only lyse eukaryotic cells but also gram-negative bacterial cells utilising a GTC buffer, while leaving tough-walled *Mtb* cells intact (4).

### Lysis and DNA extraction

Mycobacteria are known to be difficult to lyse (22). Failure to achieve adequate lysis in the context of dsWGS can result in a biased representation of target *Mtb* DNA relative to contaminating DNA (10, 22). DNA concentrations post extraction were available for four of the studies included in this review (Figure 3). DNA extraction methods varied across the reviewed studies (Supplementary Table 3). All studies, except Soundararajan *et al.* (2020) and Macedo *et al.* (2023), employed a combination of chemical and mechanical cell lysis. Soundararjan *et al*., extracted DNA utilising the Ultra Deep Microprep DNA isolation kit (Molzym, Bremen, Germany), while Macedo *et al*. utilised the QIAmp DNA Mini Kit, both of which according to the manufacturer’s instructions include both chemical and enzymatic means to facilitate lysis but omit any mechanical steps. None of the studies assessed the effect of specific pre-processing/DNA extraction steps on total DNA concentration by molecular quantification of DNA before and after any given step.

**Figure 3:**
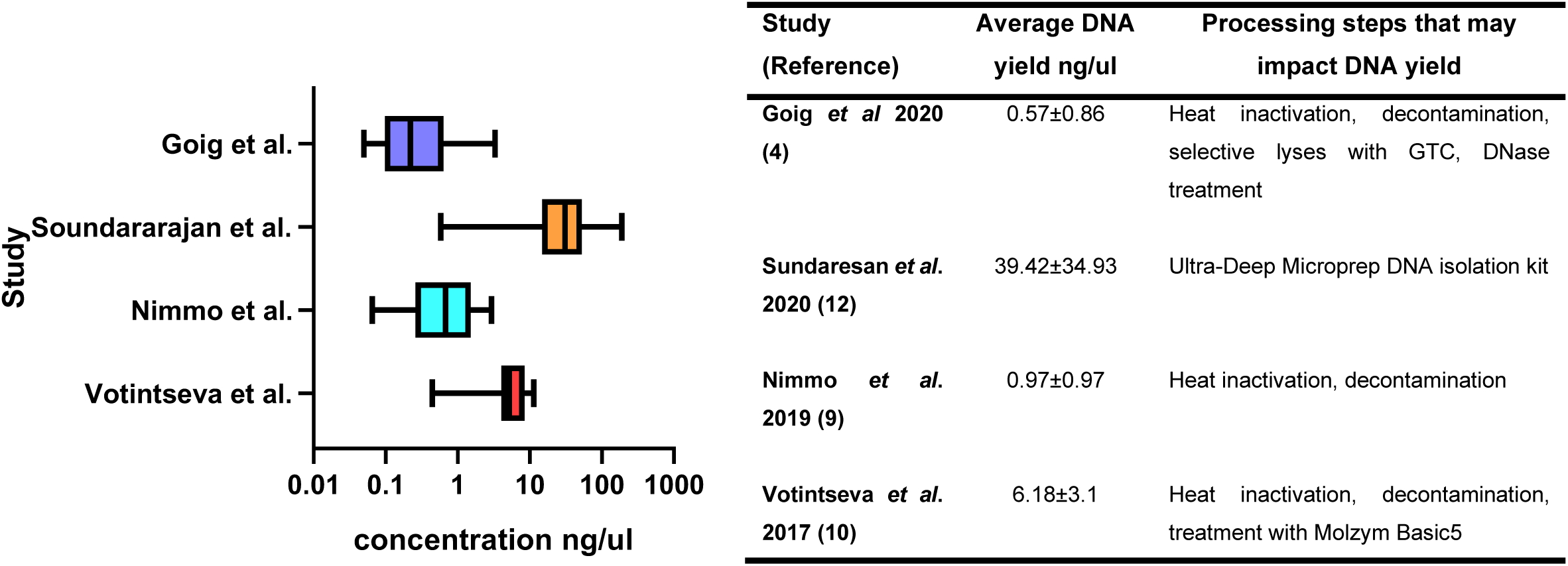
DNA concentration measured after DNA extraction and prior to target capture (if any) for the 4 studies with available data.

### Target enrichment

Six studies used DNA or RNA bait capture to enrich for *Mtb* DNA (3, 4, 9–12, 18, 21). However, only two studies (Brown *et al.* (2015) and *Goig et al.* (2020)), compared sequencing yield with or without bait capture to measure the increase of sequencing reads attributed to the *Mtb* target . The former compared the percentages of on-target reads (%OTR), and the mean sequencing depths for two sputum samples. Brown *et al.* reported a percent of on-target reads (%OTR) of 0.3%, with a sequencing depth of 4.6x without bait capture, compared to 82% and 200x respectively with bait capture. Goig *et al*., quantified the target *Mtb* DNA in the input and used this information to target bait enrichment to the lowest input samples. We used the raw data provided with this publication to assess percentage of *Mtb* target pre and post bait capture (Figure 4). The average % of *Mtb* target pre-bait capture was 1,67% compared to 48,5% post-capture (4).

**Figure 4.**
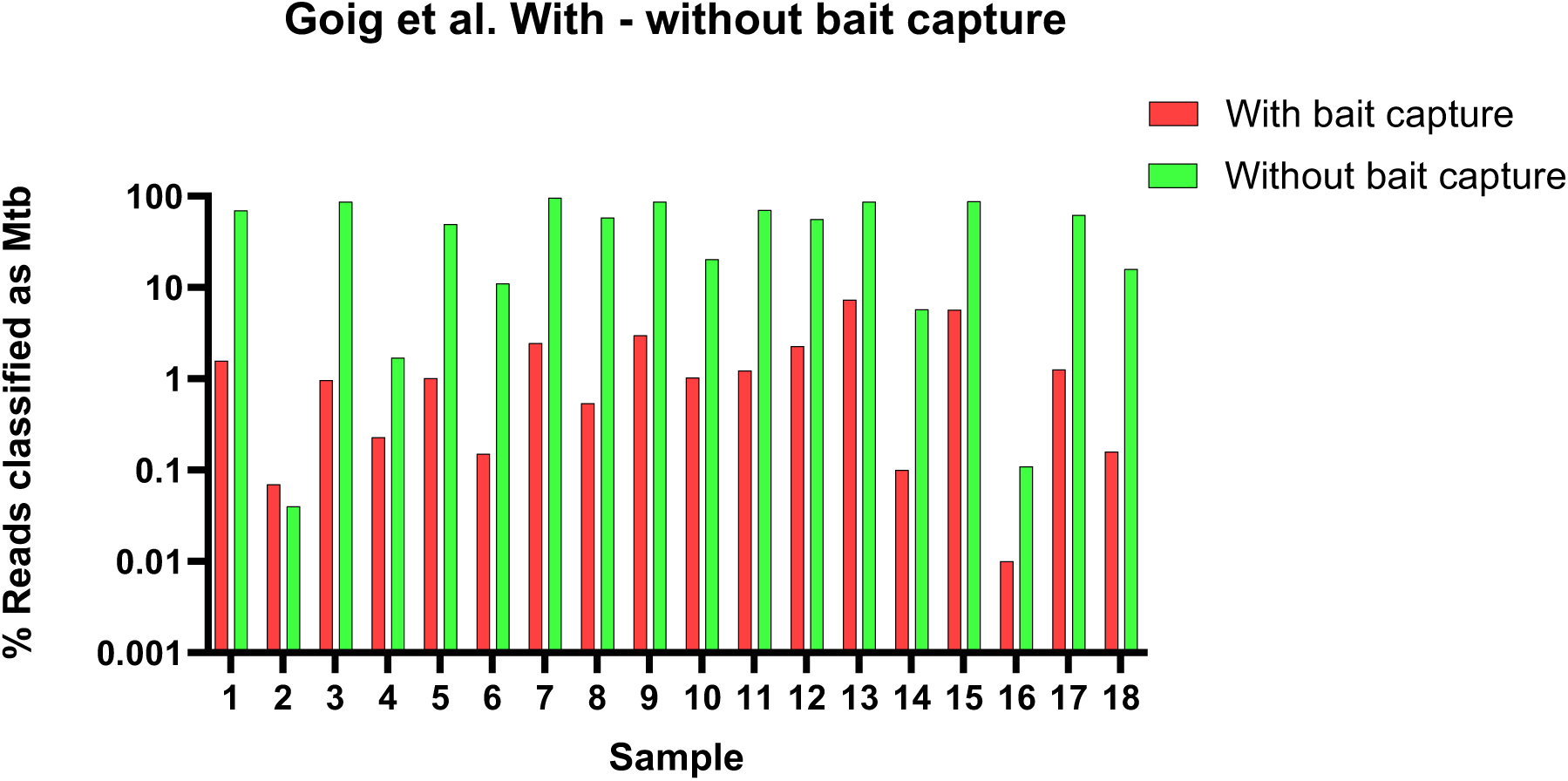
Increase in *Mtb* target read % in samples sequenced with and without bait capture from Goig *et al.,* 2020 (4). Note the y-axis is on the logarithmic scale.

### Processing steps predictive of successful dsWGS

The effects of sample characteristic (smear grade), and processing steps prior to target enrichment and sequencing (mechanical disruption for lysis, enzymatic/chemical lysis, decontamination with NaOH and heat treatment) were studied as predictors dsWGS success using logistic regression (Methods) with data extracted for 289 samples. The effect of processing steps was consistent across the dsWGS success metric used (>95% of drug resistance regions covered at >5x read depth or >90% of the whole genome covered at >5x depth, Table 3, Supplementary Table Y) and we observed no evidence of batch effects by study (Supplementary table Z). The results identify samples with higher smear grade as more likely to be successfully sequenced directly, as expected. In addition, mechanical disruption, and enzymatic/chemical lysis were also associated with higher dsWGS success prior to target capture. Sputum decontamination with NaOH on the other hand resulted in lower dsWGS success when used prior to target capture (Table 3).

**Table 3.** Sputum processing step that are significantly associated with dsWGS success based on the whole genome coverage generalized linear model.

| Characteristic | OR | 95% CI | p-value |
| --- | --- | --- | --- |
| Smear neg | ref |  |  |
| 2+,1+,scanty | 3.4 | (0.98, 12.1) | 0.07 |
| 3+ | 14.7 | (3.5, 62.1) | 0.0005 |
| Mechanical lysis | 193.3 | (11.7, 3197.8) | 0.008 |
| Enzymatic/Chemical lysis | 18.5 | (1.9, 183.1) | 0.02 |
| Decontamination | 0.005 | (0.001, 0.03) | 7e-06 |
| Heat Inactivation | 2.25 | (1.1, 4.5) | 0.02 |

Associations were performed using a sample-level logistic regression model pooled across studies with dsWGS success defined as whole genome coverage >90% at read depth of >5x per site. OR = Odds Ratio, CI = Confidence Interval.

## Discussion

This systematic review and meta-analysis summarize key experimental steps that may influence the success of dsWGS for *Mtb*. We identified and reviewed 226 study records, ultimately including 8 studies that met our inclusion criteria. Included studies varied in sample size, ranging from 34 to 100 participants/*Mtb* isolates, and utilized various approaches for the pre-process of sputum samples prior to sequencing. The current review demonstrates that although there is overlap in the applied methodology across studies there is simultaneously a lot of variability in the specific processing steps employed for sequencing directly from sputum. This review classifies key pre-processing steps that may or may not contribute to the success of dsWGS, these include sputum homogenization, decontamination, heat inactivation, depletion of host and commensal microbial DNA, lysis and DNA extraction, and target enrichment through DNA/RNA bait capture (3, 9, 11, 12, 21).

Due to the amount of contaminating cells and DNA present in sputum the main overlapping trend across studies is the utilisation of bait capture and enrichment probes which was utilised by all the reviewed studies except that of Votintseva *et al*. (2017). Bait capture relies on the hybridisation of *Mtb* specific biotinylated DNA/RNA probes that bind to complimentary *Mtb* DNA, bait/target DNA hybrids are then captured with streptavidin magnetic beads and pulled down magnetically allowing for the selective enrichment of the *Mtb* target (23). Studies by Brown *et al*. (2015) and Goig *et al*. (2020), have sequenced samples both enriched and unenriched for comparison clearly demonstrating the effectiveness of the addition of target capture and enrichment systems for dsWGS, however the impact of additional pretreatment steps has not yet been elucidated (3, 4). Homogenization, heat inactivation, and the depletion of contaminating host and bacterial cells/DNA are key steps identified during the literature review that could potentially enhance *Mtb* enrichment. Currently reviewed studies have not critically assessed the impact of various sample treatment/preparation steps on the final outcome/success of dsWGS, but have consistently highlighted correlation between smear grade/*Mtb* load and improved sequencing performance (11, 12, 21).

Other than bait capture, there is limited data on the impact of specific sputum processing steps on dsWGS success (3, 4, 9–12, 21). To address this, we used logistic regression to perform a meta-analysis of the effect of a subset of these steps on dsWGS success controlling for the *Mtb* load in the sputum as measured by smear microscopy. Study data allowed the evaluation of the effects of four processing steps and only prior to target capture: specifically, sputum decontamination with NaOH, mechanical disruption, enzymatic/chemical lysis, and heat treatment. The results support the latter three steps as significantly increasing the success of sputum dsWGS. The *Mtb* cell wall is difficult to lyse and this may explain why a combination approach which involves chemical, enzymatic and mechanical lysis contributes to improved *Mtb* DNA recovery and thus also potentially improved sequencing results (4, 24). Although heat inactivation was employed with the goal of sterilizing the sample and thereby reduce the biohazard risk of downstream processing, George *et al*. (2020) (18) demonstrated that heat inactivation can enrich for tough-to-lyse cells like *Mtb* in the setting of thermal protection buffer supporting the meta-analysis association between heat duration and temperature and dsWGS success that we observe across studies. The available data limited our ability to study the effect of other processing steps, such as host DNA depletion, using the MolYsis Basic5 kit, the Ultra Deep Microprep DNA isolation kit, or a GTC solution combined with DNase treatment (4, 10, 12). Confirming the effectiveness of these methods in depleting host DNA and enriching *Mtb* will require additional future study.

Our meta-analysis supports sputum decontamination using NaOH treatment as decreasing dsWGS success in the setting of target capture. The use of NaOH is standard practice to reduce live contaminants in sputum prior to *Mtb* culture (19). While it is known to create a highly alkaline environment that is inhospitable to most microorganisms except *Mtb*, its impact on dsWGS, specifically in terms of *Mtb* enrichment and potential loss of target cells and DNA, has not been thoroughly evaluated (25, 26). NaOH can selectively lyse contaminating cells that are not of interest for downstream analysis, but studies support a risk of *Mtb* cell loss (4, 19, 25, 26). Our finding raises a need to reevaluate the effect of NaOH treatment on sample composition and the exploration of alternative methods as potentially more suitable for WGS which does not require viable *Mtb* bacilli.

## Conclusion

Despite the observed heterogeneity of approaches for dsWGS, a common trend observed during the course of this systematic review and meta-analysis is the utilization of target capture and enrichment probes, which is observed to be highly effective in enhancing *Mtb* sequencing from direct patient samples (3, 4, 9, 12). The efficacy of these probes nevertheless depends on the overall *Mtb/Mtb* DNA load in sputum. Target capture probes are expensive and alternative or additional processing steps that can deplete contaminants or directly enrich for *Mtb* DNA will be beneficial to facilitate dsWGS and reduce cost (4, 10). Future research should thus focus on refining the identified pre-processing steps to enhance the robustness and also reliability of dsWGS with the ultimate aim of developing standardized pre-processing protocols to advancing DR profiling directly from clinical sputum samples (4, 9, 10, 12). Considering the importance of *Mtb* load highlighted in the current study a suggested additional research direction is to optimise sputum collection, standardize sputum volume, storage, handling and transport with the aim of further improving *Mtb* bacilli yield prior to the application of target capture and enrichment (27, 28).

## Supporting information

SupplementaryTables_Regression

## Acknowledgements

This work was supported by the National Institutes of Health (NIH - R01 AI155765). All claims expressed in this article are solely those of the authors and do not represent those of their affiliated organizations, or the funders. The funder had no role in study design, data, data analysis, the decision to publish, or preparation of the manuscript.

